## Supplementary figures for "Distinct contributions of the dorsal and ventral hippocampus to spatial working memory and spatial coding in the prefrontal cortex"

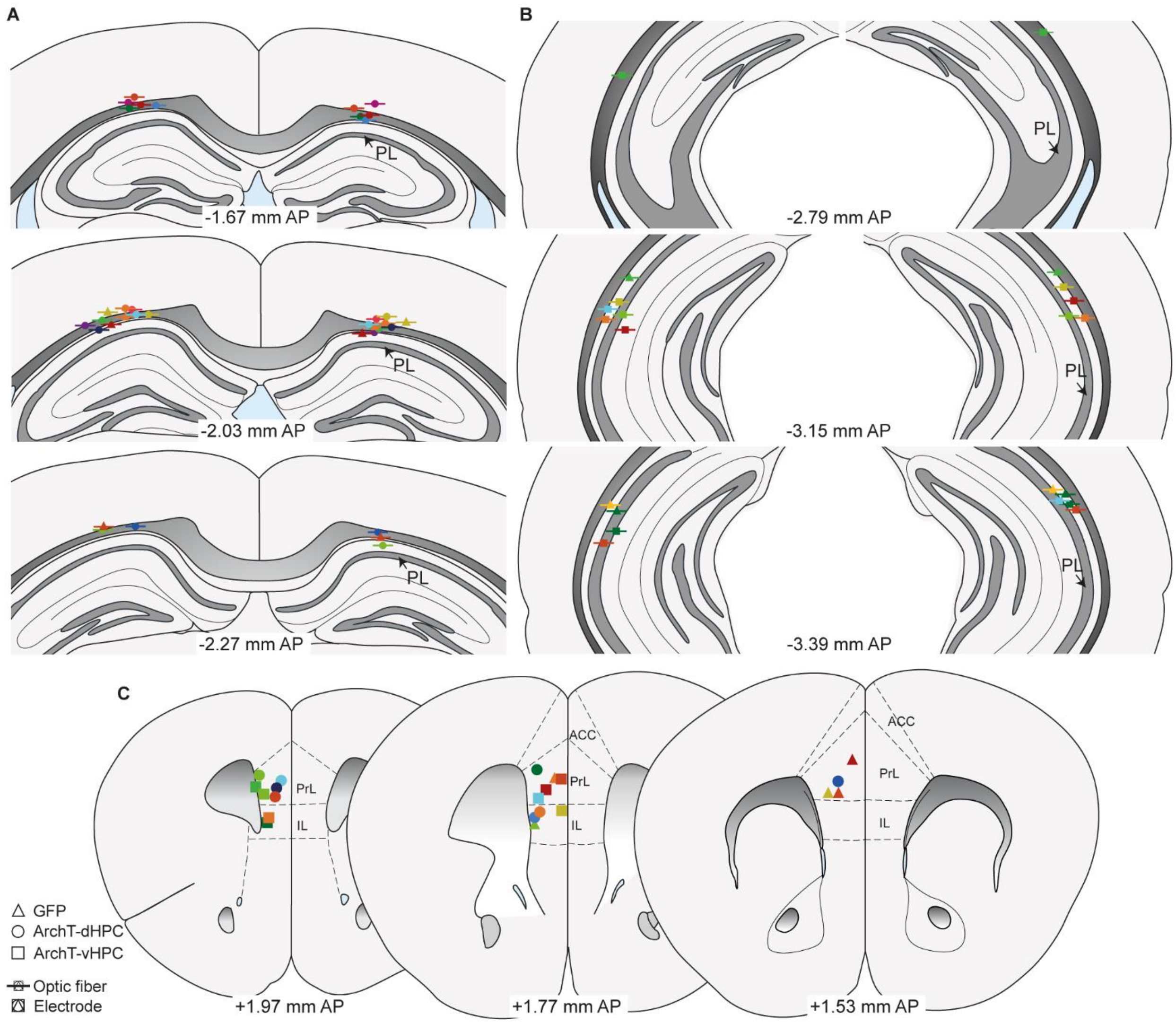


**Supplementary Figure 1: Placement of optic fibers and recording electrodes.**
**A-B**, Placement of optic fibers in dHPC (**A**) and vHPC (**B**). **C**, Positions of recording electrodes in the mPFC. Numbers indicate anteroposterior position relative to bregma. Colors indicate the placements of each animal and the symbols indicate the experimental group to which they belong. PrL, prelimbic cortex; IL, infralimbic cortex; ACC, anterior cingulate cortex. Atlas pictures are adapted from Franklin and Paxinos (2008).

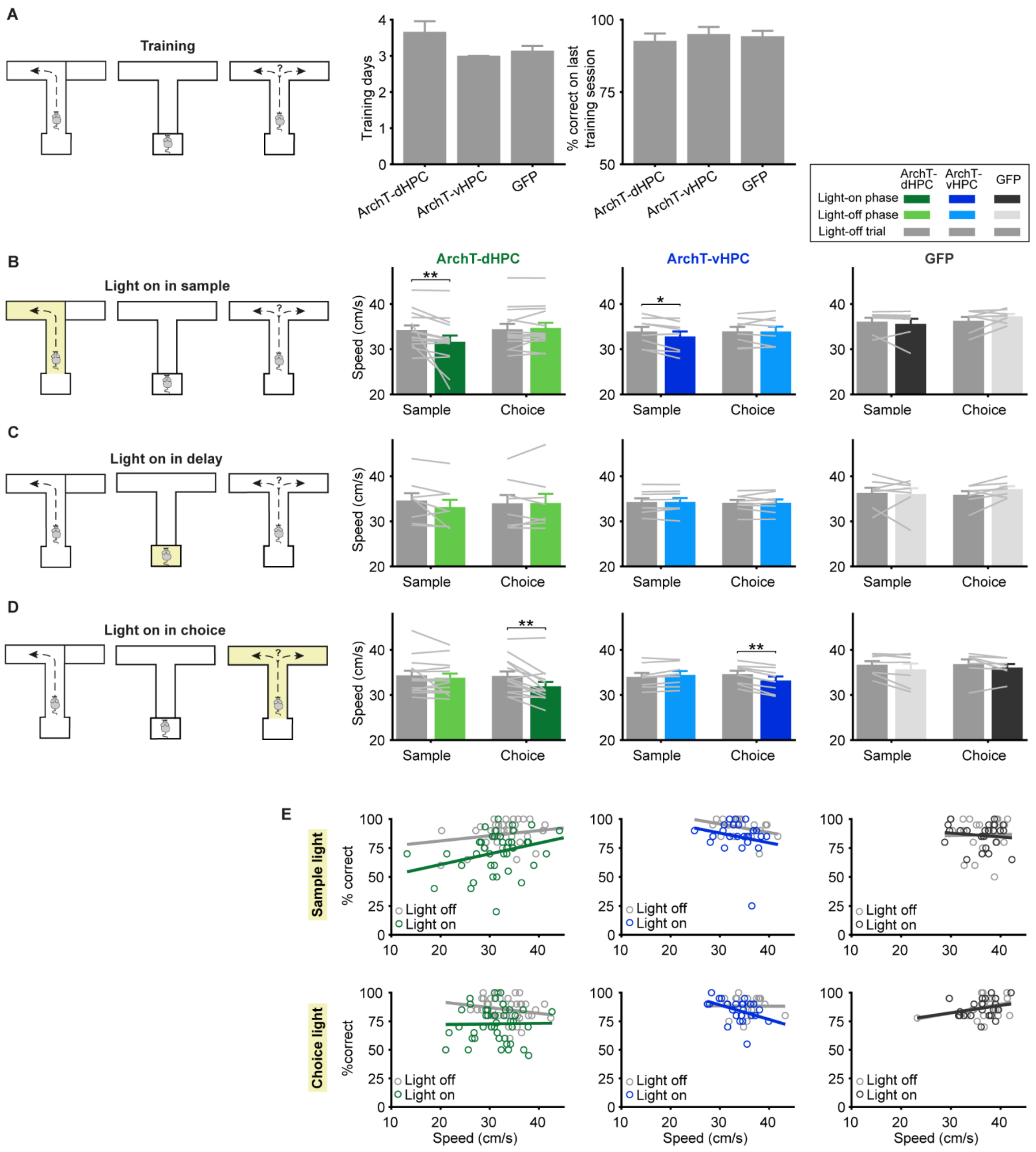


**Supplementary Figure 2: Analysis of behavioral performance and running speed.**
**A**, The number of training sessions required to reach criterion performance (left) and performance on the last training session (right) did not differ between the three experimental groups. **B-D**, Effects of hippocampal silencing during the sample phase (**B**), delay phase (**C**) and choice phase (**D**) on animals' running speed in the sample and choice phase. Dark colored bars show running speed during task phases in which light was delivered ('light-on phase'), lighter colored bars show running speed in task phases where light was not delivered ('light-off phase') and gray bars show running speed in trials without light delivery in any phase ('light-off trials'). Statistical analyses were only performed for task phases in which light was delivered (see results for more details). **E**, Relationship between running speed and performance during light on and light off trials. Each dot represents the performance and median running speed in a single session, measured in the same task phase in which light was delivered. Lines represent the linear fit of performance onto running speed for light-off and light-on trials. Error bars indicate mean ± s.e.m. across animals. *p<0.05, **p<0.01, sign-rank test.

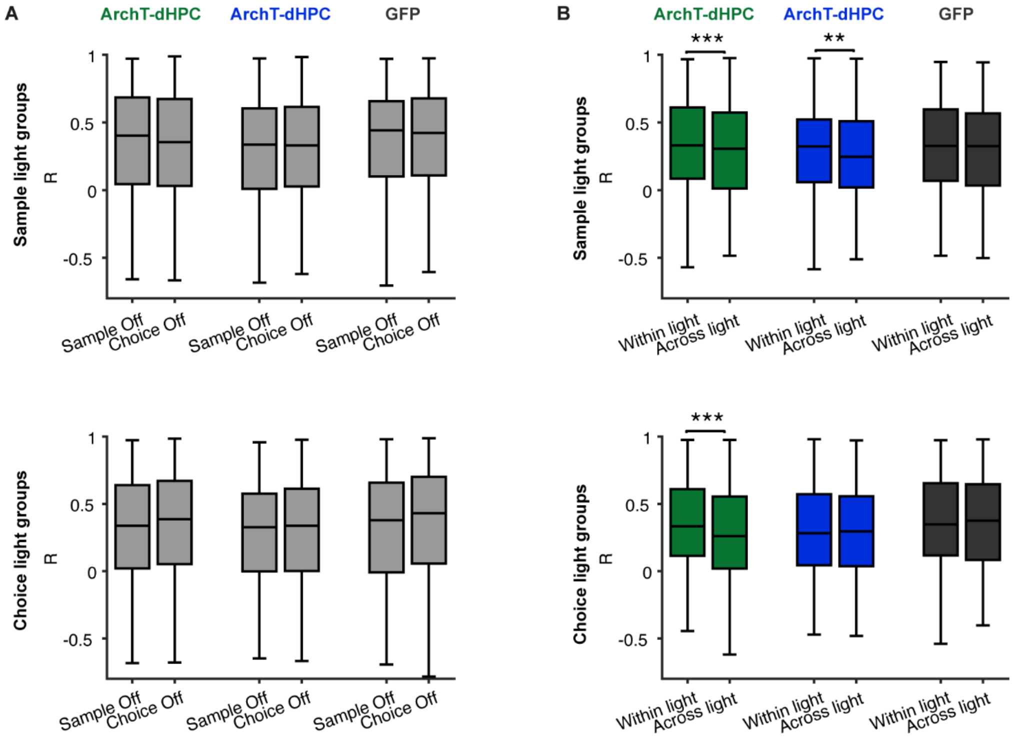


**Supplementary Figure 3: Effect of hippocampal silencing on linearized firing rates.**
**A**, Correlations between linearized firing rates in even and odd light-off trials were similar for sample ('Sample Off') and choice phases ('Choice Off'), regardless of whether light was delivered in the sample (top) or the choice (bottom) phase. **B**, For each task phase, correlations between even and odd light-off trials and between even and odd light-on trials ('Within light) were compared with correlations between even light-off and odd light-on trials and between even light-off and odd light-on trials ('Across light') for sample (top) and choice phase (bottom) silencing. These results confirm that linearized firing rates are altered by silencing of the dHPC and vHPC in the sample phase and by the dHPC in the choice phase (compare with Figure 4D). Box plots represent the median (line), 25th and 75th percentiles (box) and the whiskers extend to the minimum and maximum values within 1.5 times the interquartile range below and above the 25th and 75th percentiles, respectively. **p<0.01, ***p<0.001 sign-rank test.
